# Integration of metataxonomic datasets into microbial association networks highlights shared bacterial community dynamics in fermented vegetables

**DOI:** 10.1101/2023.11.10.566590

**Authors:** Romane Junker, Florence Valence, Michel-Yves Mistou, Stéphane Chaillou, Hélène Chiapello

**Affiliations:** Université Paris-Saclay, INRAE, MaIAGE, Jouy-en-Josas, France; INRAE, Agrocampus Ouest, STLO, Rennes, France; Université Paris-Saclay, INRAE, MICALIS, Jouy-en-Josas, France

## Abstract

The management of food fermentation is still largely based on empirical knowledge, as the dynamics of microbial communities and the underlying metabolic networks that produce safe and nutritious products remain beyond our understanding. Although these closed ecosystems contain relatively few taxa, they have not yet been thoroughly characterized with respect to how their microbial communities interact and dynamically evolve. However, with the increased availability of metataxonomic datasets on different fermented vegetables, it is now possible to gain a comprehensive understanding of the microbial relationships that structure plant fermentation.

In this study, we present a bioinformatics approach that integrates public metataxonomic 16S datasets targeting fermented vegetables. Specifically, we developed a method for exploring, comparing, and combining public 16S datasets in order to perform meta-analyses of microbiota. The workflow includes steps for searching and selecting public time-series datasets and constructing association networks of amplicon sequence variants (ASVs) based on co-abundance metrics. Networks for individual datasets are then integrated into a core network of significant associations. Microbial communities are identified based on the comparison and clustering of ASV networks using the “stochastic block model” method. When we applied this method to 10 public datasets (including a total of 931 samples), we found that it was able to shed light on the dynamics of vegetable fermentation by characterizing the processes of community succession among different bacterial assemblages.

**IMPORTANCE:** Within the growing body of research on the bacterial communities involved in the fermentation of vegetables, there is particular interest in discovering the species or consortia that drive different fermentation steps. This integrative analysis demonstrates that the reuse and integration of public microbiome datasets can provide new insights into a little-known biotope. Our most important finding is the recurrent but transient appearance, at the beginning of vegetable fermentation, of ASVs belonging to *Enterobacterales* and their associations with ASVs belonging to *Lactobacillales*. These findings could be applied in the design of new fermented products.

## INTRODUCTION

Over the last 20 years, the development of low-cost sequencing technologies has led to the creation of a large number of microbiome datasets, mainly generated using metataxonomic analyses based on 16S rRNA metabarcoding technology. For example, the number of papers using metataxonomic or metagenomic approaches to study the microbial communities of food increased six-fold between 2015 and 2021, and currently exceeds 600 [1]; similarly, within the NCBI database, the Taxonomy ID “Food metagenome” (NCBI:txid870726) is associated with 770 BioProjects. In keeping with the principles of Open Science, most of these publication-associated datasets are available in public repositories such as SRA (the Sequence Read Archive of NCBI), ENA (the European Nucleotide Archive of EBI), or DDBJ (the DNA Data Bank of Japan). To promote the reuse of certain kinds of datasets, specialized databases have been developed, such as MGNIFY for microbiome data [2]. The availability of such vast amounts of metataxonomic data provides an unprecedented opportunity to develop new integrative tools for comparing and better understanding various microbial ecosystems. However, these efforts face numerous challenges related to data reusability (e.g., data availability, metadata quality, data preprocessing) and the most appropriate ways of identifying biologically informative features in a collection of metataxonomic studies. In this work, we address these challenges by developing a method for exploring public datasets related to the microbiota of fermented vegetables and performing meta-analyses of previous research (i.e., reusing independent datasets, integrating them into a larger analysis to generate new knowledge).

Our choice of ecosystem was motivated by current interest in the bacterial communities involved in the fermentation of vegetables [3, 4, 5]. Plant-based fermented foods diversify human diets and possess interesting properties in terms of sustainability and nutritional quality. These products require little energy to produce and preserve, and their consumption confers several benefits on human health [6, 7]. With this study, we wanted to assess whether public datasets that are already available for fermented vegetables could help to improve our knowledge on the ecological dynamics taking place in these products. Fermented vegetables are created through the (usually spontaneous) activity of heterofermentative and homofermentative lactic acid bacteria (LAB) naturally present on the raw material [8]. In Europe, the most popular example of this kind of food is sauerkraut, for which the use of pre-selected starter strains remains uncommon even for large-scale production [9]. A combination of low pH and the anaerobic conditions resulting from the fermentation process are the main factors that select for the beneficial anaerobic LAB essential in the production of good-quality fermented vegetables [3]. These bacteria are a broad and diverse group of species classified in phylum *Firmicutes*, class *Bacilli*, and order *Lactobacillales*, and include representatives from the families *Lactobacillaceae*, *Streptococcaceae*, *Enterococcaceae*, *Carnobacteriaceae*, and *Aerococcaceae* [10].

It should be noted that, to date, most studies have focused on describing the microbial communities present at the end of the fermentation process [4, 5], while the dynamic succession of various microbial populations during fermentation has received little attention. This represents an important gap in knowledge, especially when compared, for example, to research on cheese microbial communities which has revealed that the proper succession of microbial populations is important to the quality of the final product [11, 12]. Two separate metataxonomic analyses that have revealed important changes in microbial dynamics during vegetable fermentation. A study on carrot juice reported a succession process involving *Enterobacteriaceae*, *Leuconostoc*, and *Lactobacillus*, while work on Suan Cai (Chinese pickles) showed that the dominant species changed from early stages of fermentation (*Leuconostoc mesenteroides*) to later ones (*Lactiplantibacillus plantarum*) Wuyts et al. [13], Yang et al. [14]. The little information that can be gathered on the subject does not allow us to identify species or consortia that might be responsible for controlling various stages of fermentation among different vegetables. In this context, the use of metataxonomic data to carry out meta-analysis could prove illuminating.

The use and comparison of amplicon data (such as the 16S-based data considered in the present work) raises certain difficulties. First, sequencing technology may vary among studies, as may the region amplified or PCR primers employed. Second, taxonomic assignment based on the 16S variable region is considered valid only to the genus level, limiting species-level interpretations [4]. There are therefore two possibilities for carrying out a comparative study of multiple datasets: comparing genus-level taxonomic profiles or comparing exact sequences, specifically, amplicon sequence variants (ASVs). The advantages of the first approach include the ability to compare different sequenced regions and to reduce the sparsity of the count matrices, while the use of ASVs enables intra-genus diversity to be taken into account [15, 16]. In both cases, the aim of this type of meta-analysis is often to identify core taxa based on criteria of abundance and prevalence [17].

The analytical design of such a study is also important. One promising approach for meta-analysis is the construction of microbial association networks, which provide additional and complementary information to classic analyses of alpha- and beta-diversity [18]. Association networks enable the identification of hub species [19, 20], taxa clusters [21], and core networks, the last of which corresponds to the intersection of several microbial association networks and can be used to identify taxa and associations shared by most networks [22]. Association networks were originally designed for macroscopic ecosystems and have only recently been adapted for the investigation of interactions within microbial assemblages [21]. They are constructed using count data from the sequenced environment, which are compositional [23], high-dimensional, and in the form of sparse matrices, thus increasing the difficulty of analysis [21]. However, compared to networks from other assemblages, the association networks in fermented ecosystems appear to be significantly smaller [16], making them easier to construct, visualize, and compare. According to Chen et al. [24], association networks can be divided into four categories, which are built using different approaches: correlation networks (CoNet [25], SparCC [26]), conditional correlation networks (SPIEC-EASI [27]), mixture networks (MixMPLN [28]), and differential networks (DCDTr). Due to the complexity of microbial interactions, all these approaches have important limitations, and no method has yet managed to capture all of the aspects of interest. Indeed, studies have even shown that classical measures such as Pearson and Spearman correlations can perform just as well as computationally time-consuming methods based on more sophisticated statistical models [29, 30].

This study presents an integrative bioinformatics approach for the meta-analysis of public amplicon datasets. The workflow includes steps designed to search for and select public time-series datasets and construct ASV association networks based on co-abundance metrics. Microbial communities are then analyzed by comparing and clustering the ASV networks. We applied this workflow to 10 publicly available datasets on the microbial assemblages of fermented vegetables. Here, we describe the value of this approach for discovering core bacterial taxa and core associations shared by different vegetables during the process of fermentation.

## RESULTS

### Design of a bioinformatics workflow for integration of metataxonomic datasets

Figure 1 depicts the main steps of the bioinformatics workflow designed to analyze and integrate the amplicon datasets. The first step involved the careful selection of public datasets focused on the microbial communities of fermented vegetables. Next, ASV count tables were constructed for each of the selected studies. Using these count tables, we then produced ASV association networks for each study that were based on four sensitive and computationally efficient metrics: Jaccard distance, Pearson and Spearman correlations between relative abundances, and a proportionality measure calculated from clr-transformed abundances. The purpose of the networks was to help visualize how microbial communities interact and evolve dynamically. Finally, the various networks were integrated together. A core network was constructed that identified which bacterial ASVs were common to most fermentations and which associations between ASVs were significantly shared among networks. In addition, a multiple SBM clustering method was used to identify a set of ASVs that were associated with each other across the different networks.

**Figure 1:**
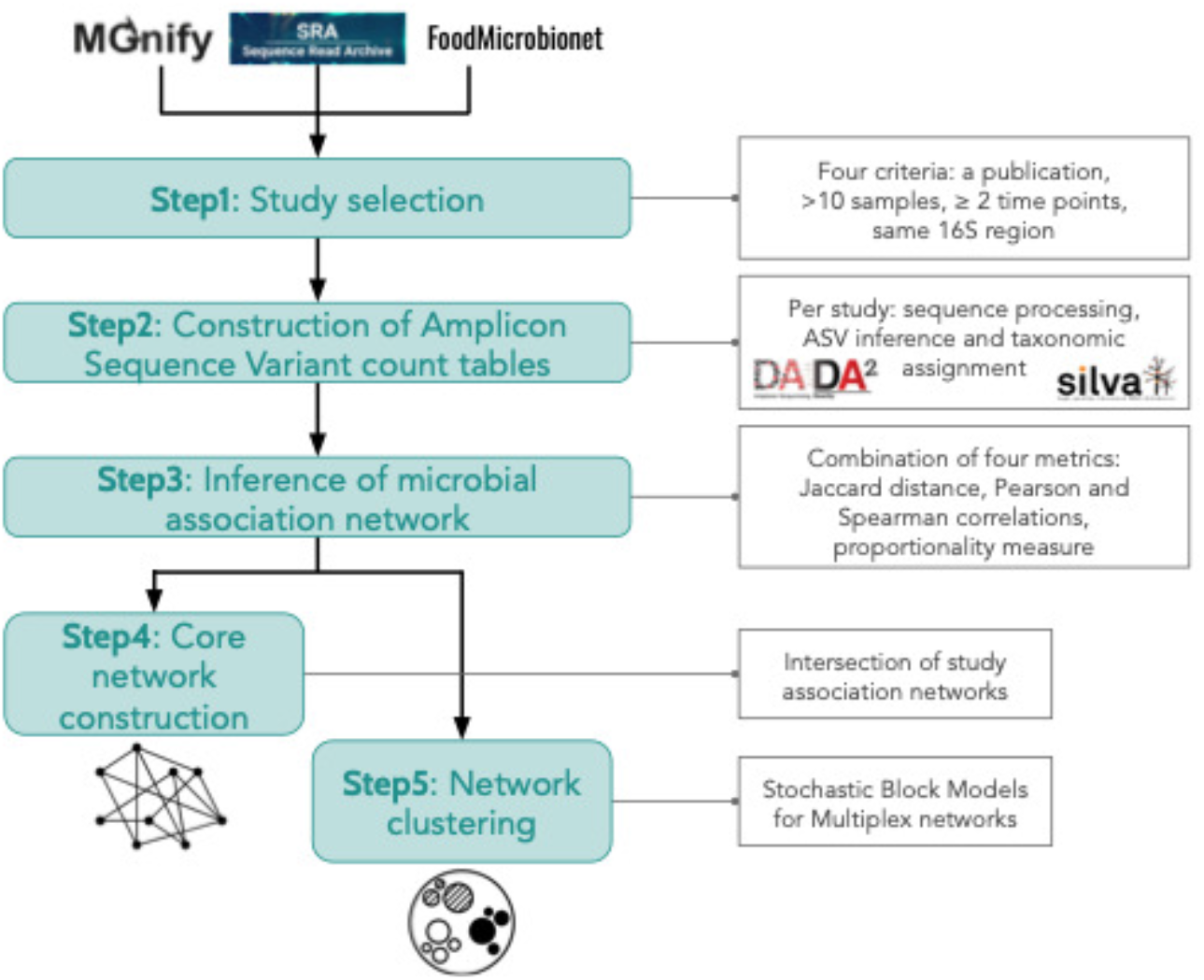
Meta-analysis approach for integrating amplicon datasets into microbial association networks to compare microbial communities of fermented vegetables.

### Selection of metataxonomic studies on fermented vegetables

Ten datasets meeting our selection criteria (see Materials and Methods section) were obtained out of 1443 studies from SRA (NCBI), 10 studies from MGnify (ENA), and 3 studies from FoodMicrobioNet. All datasets contained sequences of the V3–V4 or V4 hypervariable region, enabling ASV comparison. The selected datasets originated from studies on five different varieties of vegetables (cucumber, carrot, cabbage, pepper, radish, used alone or in a mixture) and comprised between 18 and 310 samples each, for a total of 931 samples (Table 1). The time scales that were examined varied among studies, as the datasets included between 2 and 12 time points. Depending on the study in question, monitoring began between 0 and 30 days after the beginning of fermentation and ended between 3 and 720 days after. All studies were conducted on spontaneous fermentations, with the exception of PRJNA751723 and PRJNA662831, which included samples from spontaneous fermentations as well as samples inoculated with various LAB (*Latilactobacillus curvatus*, *Leuconostoc gelidum*, *Latilactobacillus sakei*, or *Weissella koreensis*). Dataset PRJEB15657 contained data from two sets of experiments (samples from a laboratory experiment and samples from a citizen science experiment), which we divided into two subsets.

**Table 1:**
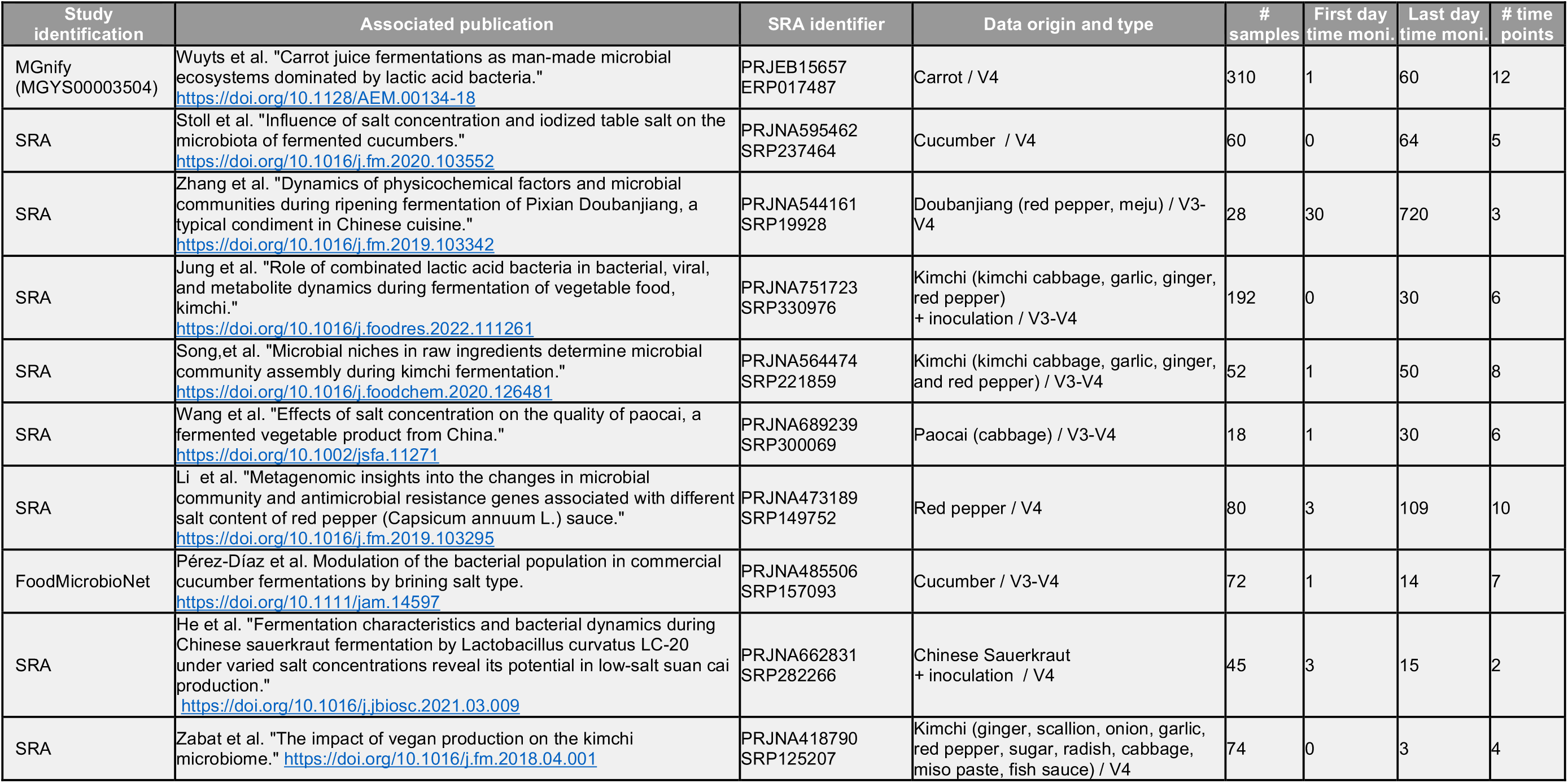
Publicly available metataxonomic studies on fermented vegetables used in the present work. . The table describes the main features of the 10 public metataxonomic studies retained for this analysis. The <study identification= column indicates the source that allowed identification of the dataset. The <associated publication= column indicates the first author, title, and DOI of the article that was used to complete the dataset metadata. The <SRA identifier= column indicates the Bioproject SRA study identifier that was used to download the corresponding public datasets. The <data origin and type= column indicates the vegetable(s) of origin (with possible indication of additional inoculation, if applicable) of the dataset and the sequenced 16S region. The last four columns indicate, respectively, the number of samples in the study, the first and last day of monitoring, and the total number of time points assessed.

### Visualization of microbial succession during fermentation through the construction of association networks

Historically, bar graphs have been used to visualize changes in the taxonomic composition of bacterial communities between samples. However, this method does not reflect the evolution of ASV associations over time. Microbial association networks, on the other hand, highlight these temporal taxonomic associations and can visually present information that is complementary to bar graphs. For each of the 10 datasets, we built association networks, of which one is presented in Figures 2A and 2B (study PRJNA689239, paocai fermentation over 30 days, captured at six timepoints). This network appeared to be composed of two subnetworks: one containing a high diversity of ASVs (including *Pseudomonadales* and *Enterobacterales*) with a weighted mean age (WMA) between 0 and 10 days, and the other containing a lower diversity of ASVs belonging to *Enterobacterales* and *Lactobacillales*, with a higher WMA (between 8 and 30 days). These observations suggest that there is a shift during fermentation from a broad initial diversity of ASVs to an assemblage dominated by LAB. Interestingly, we observed the same patterns in the PRJNA564474 study (Fig. 2D; kimchi fermentation over 50 days and eight timepoints). However, a notable difference from the paocai study was that the first subnetwork was present at WMAs ranging from 0 to 50 days, and the second, composed only of *Lactobacillales* ASVs, appeared at 10 to 50 days. This structure suggests that some of the samples failed to ferment, as observed for sample SRR10127549 in the bar graph.

**Figure 2:**
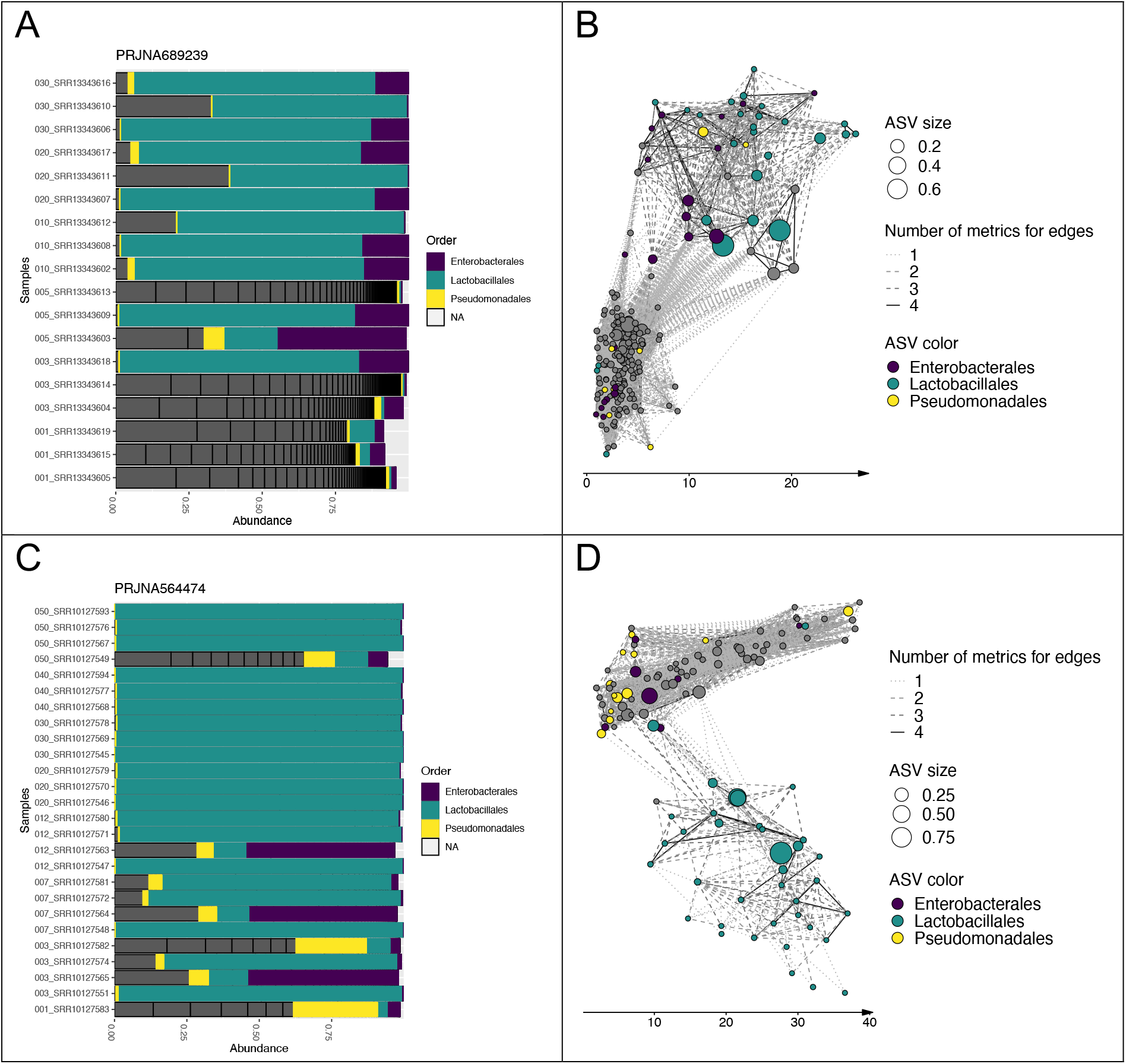
Microbial association networks for studies PRJNA689239 and PRJNA564474 highlight the dynamic evolution of microbial communities during fermentation. (A) and (C): Barplots depicting relative abundances in each sample for studies PRJNA689239 and PRJNA564474, respectively. Samples are ordered by age. Gray color indicates a taxonomic order other than *Enterobacterales*, *Lactobacillales*, and *Pseudomonadales*. (B) and (D): ASV association networks for studies PRJNA689239 and PRJNA564474, respectively. Each node represents an ASV; node size reflects its maximum relative abundance and color represents its taxonomic order. The x-axis corresponds to the weighted mean age (WMA) of the samples in which the ASV was detected, measured in days, and weighted by ASV relative abundance. An edge between two nodes indicates an association that was detected according to at least one metric.

A similar network pattern was observed for 8 of the 11 networks analyzed (Fig. S1). The overall pattern could be described as follows: samples initially contained a high diversity of ASVs (featuring *Pseudomonadales* in particular) with a low WMA; then, as the WMA increased, nodes corresponding to ASVs from *Enterobacterales* and then *Lactobacillales* appeared, with numerous associations between them. However, we would like to emphasize a few points to keep in mind when interpreting these networks. The WMA of an ASV does not reflect the exact time point at which the ASV first appears. Indeed, during each of the vegetable fermentations, all ASVs were present from the beginning of the fermentation process. This measurement may also represent both living and dead bacterial populations because the DNA of dead bacteria may be recovered and sequenced as well. Hence, the use of WMA to organize an association network merely provides a general picture of the temporal dynamics of ASVs over a fermentation process, highlighting the main “peaks” of presence and potential species associations.

We also analyzed the three networks that did not exhibit this succession of communities (PRJNA473189, PRJNA662831, PRJNA544161; see Fig. S1). A common feature of these three studies was a shift in timing compared to the others: more precisely, sampling did not start until three days after the onset of fermentation. Therefore, it is possible that the successional shift in microbial communities took place before the first sampling point. This hypothesis is supported by the observation that the taxonomic profile of the pepper and sauerkraut samples (PRJNA473189 and PRJNA662831) did not change over time. In the case of doubanjiang (PRJNA544161), a fermented product containing numerous ingredients (beans, soya, rice, spices), ASVs belonging to *Enterobacterales* appeared to proliferate relatively late, as observed on the bar graph (Fig. S1).

### Comparison of association networks to identify a core network of bacterial communities

To integrate the 11 association networks, we constructed a core network, i.e., the intersection of several networks (Fig. 3). In Figure 3A, it can be seen that the 11 networks shared 3 vertices (ASVs) overall, and pairwise analyses revealed between 10 and 58 vertices that were shared between a given pair of networks. Similarly, pairwise analyses detected between 3 and 296 edges that were shared by two networks, but no edges were shared by more than nine networks (Fig. 3B). To evaluate the statistical significance of the edge intersections, we compared them with a null model using a Kolmogorov-Smirnov test; the results rejected the null hypothesis that our set of networks followed the same distribution as the null model for intersections between two, three, four, five, or six networks (p-value < 0.05 for 100 cases). This means that those network subsets share associations in a significant way.

**Figure 3:**
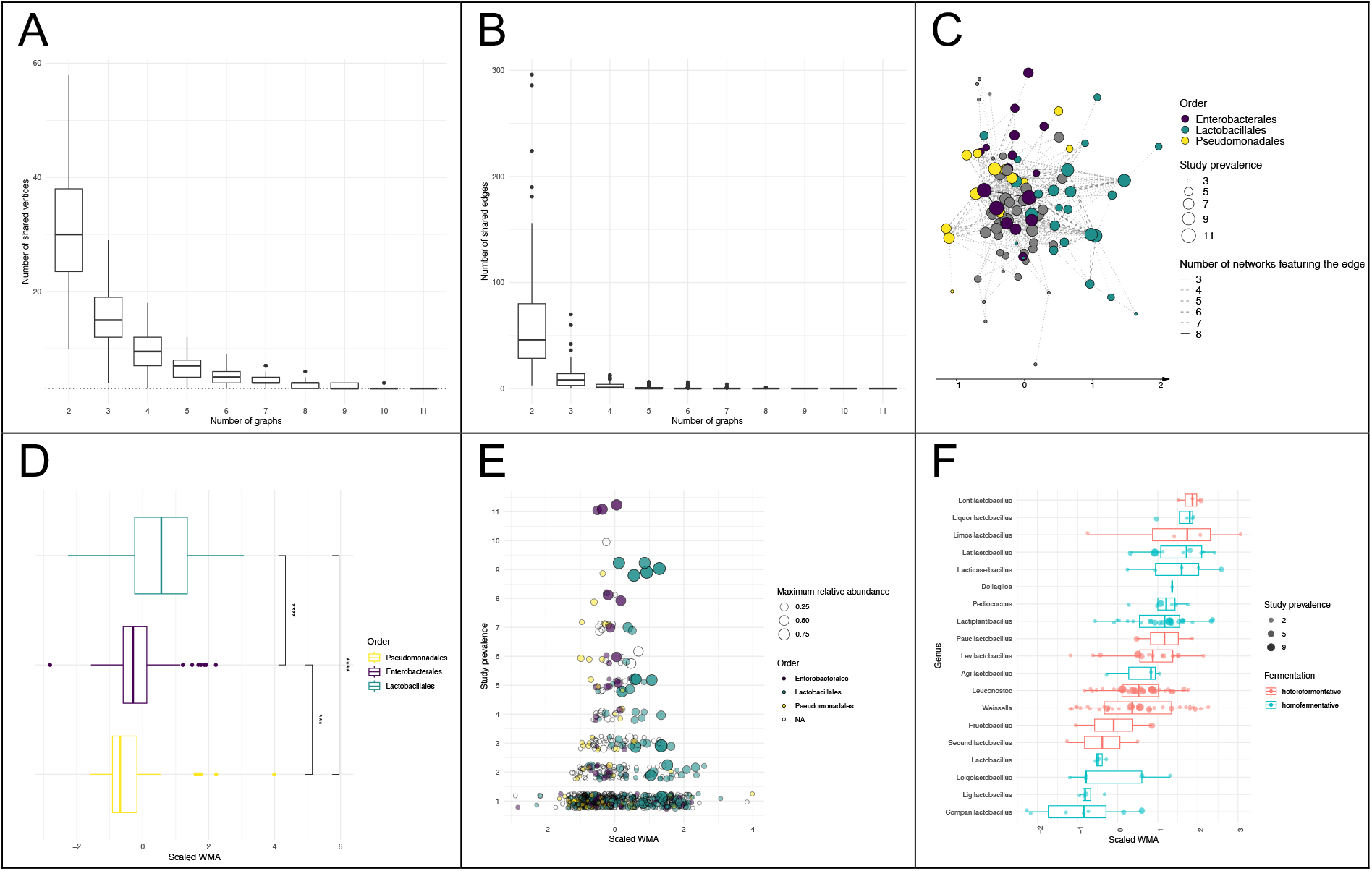
Core network and succession of bacterial communities. (A) Boxplot showing the number of vertices in the core networks built from the intersection of 2 to 11 networks. The dotted gray line corresponds to three vertices. (B) Boxplot showing the number of edges in the core networks built from the intersection of 2 to 11 networks (C) Core network built from ASV associations found in at least three networks. The line type of an edge represents the number of times the ASV association was found. The node position on the x-axis is the mean scaled WMA. ASVs are colored by taxonomic order. (D) Boxplot showing the differences in mean scaled WMA between ASVs affiliated with the orders *Pseudomonadales*, *Enterobacterales*, and *Lactobacillales*. (E) Scatterplot of ASVs colored by taxonomic order, depicting their prevalence in relation to mean scaled WMA. (F) Boxplot showing the differences in mean scaled WMA among genera in family *Lactobacillaceae*. Each dot is an ASV.

We then constructed core networks based on the intersections between two to six networks (all shown in Fig. S2). The core network built using microbial associations present in at least three networks (Fig. 3C) included 97 ASVs (out of a total of 975 used to construct the 11 networks). Among them, 13 were affiliated with order *Pseudomonadales*, 17 with *Enterobacterales*, and 25 with *Lactobacillales*. In representing the core network, we used the scaled WMA on the x-axis. The rationale of the scaled WMA was to normalize time data and to establish a common time scale between the various studies. Indeed, the WMAs are not directly comparable between studies because the time points measured varied from one study to another.

Analysis of the different significant core networks revealed that, despite all of the differences between experiments (type of sequencing, fermentation conditions, time scale), there appeared to be a common temporal structure in the microbial dynamics of fermented vegetables. In particular, after a mean scaled WMA of 0.5, *Lactobacillales* ASVs tended to predominate. Furthermore, we also observed a shift from the initial microbial population of vegetables to one dominated by *Enterobacterales*, and then a second shift to *Lactobacillales*.

This observation was confirmed by a clear difference in scaled WMA among all ASVs corresponding to *Pseudomonadales*, *Enterobacterales*, and *Lactobacillales*, as shown in Figure 3D. Figure 3E highlights this trend and also shows that the ASVs with the lowest and highest scaled WMAs were less often shared among studies (less than three graphs when WMA was lower than −1 or higher than 2) than those with median WMA values. This suggests that the initial flora, as well as the LAB present mainly at the end of fermentation, tended to be more specific to a given experiment than other ASVs. Moreover, the ASVs belonging to *Enterobacterales* were more likely to be shared between networks than those corresponding to *Lactobacillales* (non-parametric Wilcoxon-Mann-Whitney test, p-value = 0.02). In fact, of the three ASVs that were detected in all experiments, all belonged to the *Enterobacterales* (*Klebsiella*, *Pectobacterium*, and an unidentified *Enterobacterales*).

Finally, we investigated distinctions between different genera within family *Lactobacillaceae* (following the new taxonomy of Zheng et al. [31]) based on the type of fermentation performed. In the core network, ASVs belonging to genera that perform hetero-lactic fermentation were more numerous than those belonging to genera that perform homo-lactic fermentation. Moreover, most members of the *Lactobacillales* were found in only one graph (143 out of 208, i.e., 69%), and among those shared by more than two graphs, 18 perform heterofermentation and 7 perform homofermentation. We can therefore conclude that LABs are generally highly specific to a fermentation process, and the ASVs that are shared among different processes are mostly heterofermentative. There was no significant difference between the scaled WMA of heterofermentative and homofermentative genera, but we did detect some expected successional shifts in genera (Fig. 3F: *Leuconostoc* and *Lactiplantibacillus*, p-value = 0.02).

### Multiple clustering to identify putative bacterial consortia shared among studies

To identify sets of ASVs that were connected in similar ways across the 11 microbial association networks, we applied the multiplex stochastic block model (SBM) graph clustering method. Ten different clusters were identified, which varied in their size and the prevalence and taxonomy of their member ASVs. All ASVs within a cluster shared similar intra-cluster and inter-cluster connection patterns. Clusters 1 to 5 contained few ASVs (between 5 and 45) that were shared between two or more networks, while clusters 6 to 10 contained many ASVs (between 94 and 463) that were mainly specific to one network (Fig. 4A). ASVs affiliated with *Lactobacillales* predominantly belonged to clusters 5, 9, and 10; this last group contained most of the *Lactobacillales* ASVs and those corresponding to the diverse initial microflora, i.e., those that were specific to each experiment.

**Figure 4:**
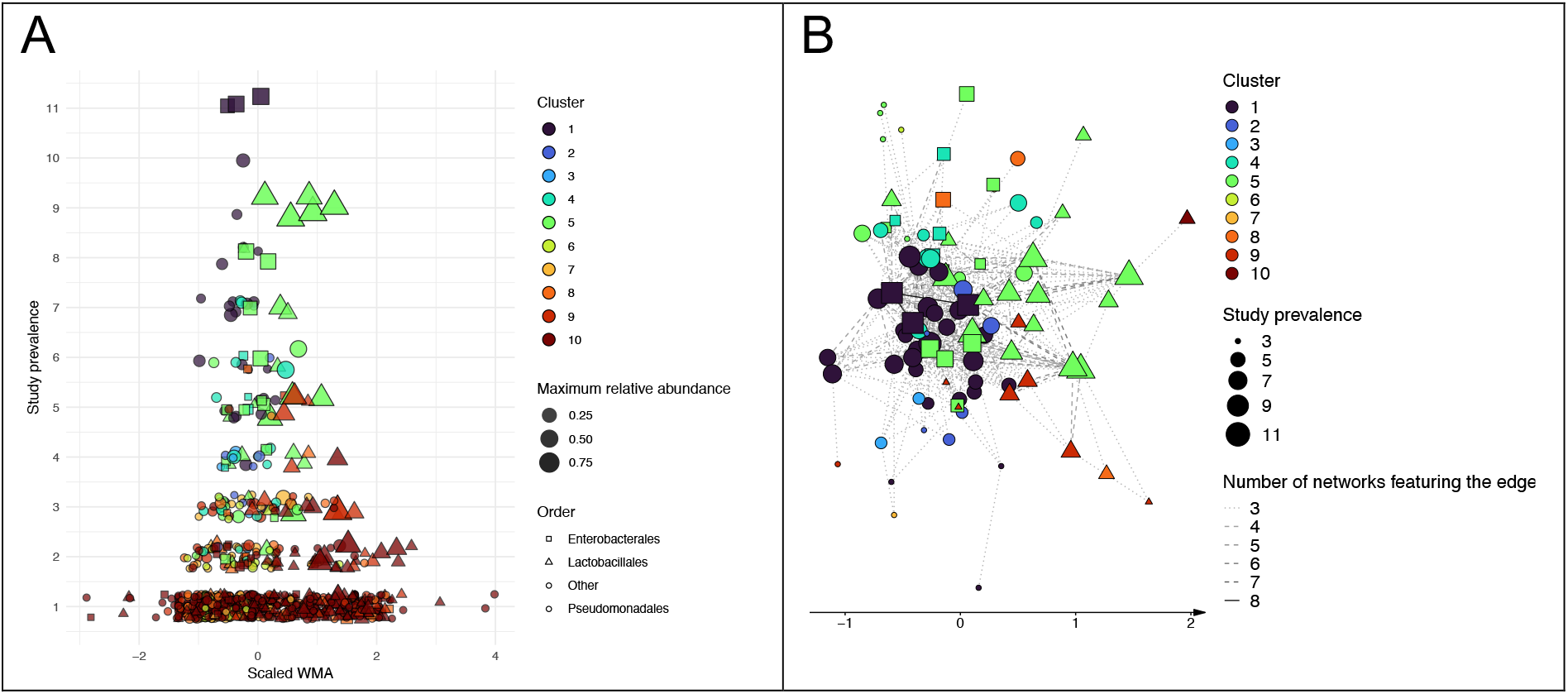
Network clustering shows an association between highly prevalent ASVs from orders. *Lactobacillales* **and** *Enterobacterales*. (A) Scatterplot of ASVs colored by cluster. The shape corresponds to the taxonomic order. (B) Core network built from ASV associations found in at least three networks, with ASVs colored by cluster. The shape corresponds to the taxonomic order.

Among the different clusters, clusters 1 and 5 were particularly interesting, as they included the majority of ASVs that were shared by more than five networks and they were the predominant clusters in the core network (Fig. 4B). Moreover, ASVs in the two clusters differed significantly in their scaled WMA (p-value = 0.013). It is possible that the ASVs in these two clusters correspond to successive bacterial communities that are common in vegetable fermentation: cluster 1 appeared to be highly associated with ASVs in the orders *Pseudomonadales* and *Enterobacterales*, while cluster 5 represented an assemblage of *Enterobacterales* and *Lactobacillales*. This community of conserved ASVs could potentially represent a shared core-consortia of early fermentation; its detailed composition is shown in Table S1.

## DISCUSSION

This work presents an integrative bioinformatics approach that utilizes association networks to combine different sets of publicly available data on the microbial dynamics of fermentation in vegetables. By combining ASV networks from different studies, we obtained valuable insights into bacterial community structure during different phases of fermentation. Historically, association networks have been used to detect potential inter-species interactions; here, we adapted this strategy to identify and visualize ASVs with similar temporal dynamics. To our knowledge, this work is the first to construct a core network representing the fermentation of different vegetables based on sequence data from multiple independent datasets. By integrating several public datasets together, we were able to characterize two successional shifts that were conserved among different fermentation ecosystems: the first from the initial microbial population of vegetables to *Enterobacterales*, and the second to an assemblage dominated by *Lactobacillales*. To test the significance of the core network we obtained, we used an approach based on comparison to a null model, which was similar to that developed by Röttjers et al. [22], with a sampling of random graphs similar to Doane et al. [32]. Indeed, the identification of core networks is a more challenging task than computation of the global intersection network [21]. With these tests, we determined that some intersections between networks would not be expected by random chance, and thus that some edges may correspond to genuine ASV dynamics shared among several studies. Finally, we complemented this approach by using the SBM method for ASV clustering, which is a technique applicable to multiplexes (a type of multi-layer network) that does not require any a priori assumptions regarding connectivity patterns. The SBM model has been used for community detection in various fields, such as sociology. More recently, it has been applied to taxonomic profiling of the human microbiome in order to uncover patterns of community structure. Specifically, it was used as a bipartite model for clustering samples and taxa [33]. In another study, the simple SBM enabled the detection of OTU clusters based on their connectivity patterns in a co-occurrence network [34]. In the present work, we applied the multiplex version of this model to a collection of networks in order to identify clusters of ASVs that share similar patterns of associations across the different networks. We were able to identify 10 clusters of ASVs, which could be used to guide the exploration and delineation of new bacterial consortia in fermented vegetables [35].

With respect to the microbial ecology of fermented vegetables, our most important finding was the recurring and transient appearance, at the beginning of fermentation, of ASVs belonging to *Enterobacterales* and their association with ASVs affiliated with *Lactobacillales*. This raises the question of their ecological function in vegetable fermentation and their impact on the properties of the final product. The hypothesis of bacterial succession in vegetable fermentation, from *Enterobacteria* to heterolactic and homolactic acid bacteria, is not entirely new. However, due to the small number of studies carried out on the subject and the extensive variability in the methodologies used, most reports have not generated convincing conclusions on the impact of *Enterobacterales* and their possible interactions with LAB. Nevertheless, based on the existing literature, several hypotheses can be put forward. *Enterobacterales* may have fermentative properties, or they may participate in nutritional mutualism that is beneficial to the development of LAB. Indeed, certain trophic relationships between LAB and *Enterobacteriaceae* have already been described. For example, some LAB generate metabolic energy using an agmatine deiminase pathway that relies on agmatine produced by *Enterobacteriaceae* [36]. In the wet coffee fermentation process, the first phase involves interactions between *Enterobacteriaceae* (with pectinolytic activity), acetic acid bacteria, and some yeasts [37]. *Enterobacteriaceae* have also been found in two other studies on fermented vegetables [38, 39], of which the former hypothesizes that the presence of *Erwinia sp.* may reflect its ability to invade compromised plant tissues or its potential ability to ferment sugar.

The meta-analysis we designed is particularly well-suited to fermented vegetable ecosystems: since these ecosystems are closed, contain relatively few taxa, and undergo a temporal succession of communities, the representation of ASV association networks is fairly easy to visualize and interpret. This approach could be easily applied to amplicon or shotgun metagenomic data for other fermented foods characterized by closed ecosystems with community shifts. One limitation of the present meta-analysis is that it was carried out on a relatively small scale (on 10 independent datasets including a total of 931 samples), due to the small number of reusable public metabarcoding datasets on fermented vegetables. This is mainly due to difficulties in accessing raw data (some samples are missing, some data are pre-processed, etc.) and metadata (sometimes incomplete and inconsistent, with manual extract from paper required). Indeed, these limitations were highlighted in a recent article [40], which recommended that data be deposited in public repositories together with assay metadata (technical features of the experiment) and biological metadata (environmental conditions of the biosamples). This, along with the adoption of other best practices, will enable wider reuse and integration of microbiome datasets on a broader scale.

This study is based on 16S metataxonomic data, more specifically, the V4 hypervariable region because it was used in the majority of the datasets found. This region is the most frequent target of studies focused on food ecosystems, along with the V3–V4 region of the 16S rRNA gene [1]. Unfortunately, this gene region has poor discriminatory power; it is able to provide reliable taxonomic assignment at the genus-level only and cannot be used to study species-level diversity (unlike, for instance, the V1–V3 region [41]). Therefore, although it is interesting to discover ASVs that are shared between different studies, this approach is ill-suited for characterizing the species- and strain-level diversity of *Lactobacillales* and *Enterobacterales*. Furthermore, the read count tables obtained for the different studies can be shaped by many biases, including differences in sample collection and storage, DNA extraction method and primer choice, variation in the number of rRNA operons [41, 42], amplification of extracellular DNA, and errors in taxonomic affiliations. Therefore, the results of any individual ASV count table must be interpreted cautiously. However, in the context of our study, the use of ASVs enabled direct comparison of sequences between studies and reduced the influence of taxonomic misclassifications [43, 44]. In addition, integrating ASVs into association networks allowed comparisons of similar dynamics between ASVs in different studies, and limited the biases that might arise from direct comparison of relative abundances.

This work demonstrates the effectiveness of using association networks for temporal meta-analysis. The approach we developed could easily be applied to new datasets or extended to incorporate new tools for association network inference, core network detection, and clustering. In the future, it could be interesting to integrate additional sample metadata (such as temperature, lactic acid concentration, pH, and/or salinity) if they were available in a standardized format and could be easily integrated to an association network. This approach could lead to the design of ideal consortia that could make vegetable fermentation safer [45] (Capozzi et al., 2017), more reproducible, and exploitable on a large scale [46].

Finally, the taxonomic profile inferred from 16S rRNA is not able to provide insights into the functional profile of bacterial communities or into the part(s) played by other microorganisms (even if their presence is minor, e.g., less than 5% relative abundance for fungi and *Archaea* in brine food according to Leech et al. [4]). Ultimately, there is a need for complementary functional studies (shotgun metagenomics, metatranscriptomics) to improve our understanding of vegetable fermentation and assess the functional interactions taking place during this process.

## MATERIALS AND METHODS

### Study selection

Datasets were obtained from three repositories: the MGnify database (on microbiome data), the FoodMicrobioNet database (on food ecosystems), and the NCBI SRA database. Studies focused on the microbial ecosystems in fermented vegetables were identified in MGnify by selecting the biome “Food production” and filtering with the term “Fermented vegetables”, while in FoodMicrobionet, we used the spoilage filter “Fermented” from studies labeled with “Vegetables and vegetable products”. From NCBI/SRA, we retrieved studies with the Taxonomy IDs “Food metagenome” (870726), “Fermentation metagenome” (1326787), and “Food fermentation metagenome” (1154581). Of the resulting studies, the only ones that were considered were those whose “SRA Run Selector” metadata contained the words “day”, “week”, “month”, “hour,” or “time”, and that had an associated publication on fermented vegetables.

We included only studies that examined at least two time points, contained more than 10 samples, and were associated with a publication (to ensure access to extensive metadata). Finally, we retained only studies that sequenced the V4 or V3–V4 hypervariable region of the 16S rRNA gene to permit comparisons of ASVs. Raw sequencing data of the resulting selected studies were retrieved from the NCBI SRA repository using home-made scripts.

### Construction of ASV count tables

Sequencing data from each study were processed using the dada2 pipeline [43] for read quality control, read filtering and trimming (with parameters truncLen = 240 or 220 depending on read length, maxN = 0, maxEE = 2, truncQ = 2), error rate learning and ASV inference, paired read assembly (with parameter minOverlap = 3), chimera removal, and taxonomic assignment to kingdom, phylum, class, order, family, and genus (using Silva database nr 99 v 138. 1). For the five studies in which the V3–V4 region of the 16S rRNA gene was sequenced, only the V4 region was retained. The ASV count table for each sample, the ASV taxonomy table, and the sample metadata were combined into one phyloseq object [47] for each study. ASVs matching mitochondrial or chloroplast DNA and samples from negative fermentation controls were excluded from the count tables.

### Inference of microbial association networks

For each study a count table was filtered to create a microbial association network. Only non-control samples with more than 15,000 reads and ASVs found in at least three samples and with an average relative abundance greater than 1e-5 were included. We chose association metrics that take into account co-presence, with Jaccard distance, as well as co-abundance, with Pearson and Spearman correlations based on relative abundances. The proportionality measure proposed by Lovell et al. [48] (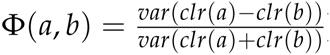) was also used following centered log-ratio transformation, performed using the function aldex.clr from the package ALDEx2 Quinn et al. [49]. Edges were traced if at least one of these four measures reached a non-stringent threshold (0.4 for Jaccard distance and 0.5 for the three other measures). The thickness of each edge reflected the number of combined metrics supporting it. A force-driven algorithm (Fruchterman-Reingold) was used to calculate the layout of each association network. This layout was preserved on the y-axis, but the x-axis was modified: the position of each ASV was the mean age of the samples in which the ASV was present, weighted by its relative abundance (hereafter named WMA for weighted mean age).

### Core network construction

The core network was constructed based on the intersections of the independent association networks created for each study. To account for the different sampling time points and fermentation rates among studies, the x-axis position of each ASV in the core network corresponded to the average of its centered and scaled positions in the original networks. A null-model statistical test was used to assess the significance of the core networks constructed from edges shared by a subset of networks or by all networks. First, we generated 100 sets of networks with the same nodes as the networks of interest but with random edges, using the “rewire” function of the igraph R package with prob = 1. Next, the distribution of edges shared by a given subset of networks or by all networks was compared between each null model and the studied set of networks with a Kolmogorov-Smirnov test.

### SBM multiplex clustering

Multiplex networks refer to a collection of networks involving the same sets of nodes but originating from different types of relationships. Here, each network corresponded to a specific study and each node corresponded to an ASV. SBM clustering was applied to multiplex networks to assign each ASV to a community (or block) according to its connection patterns. The estimateMultiplexSBM function from the R package sbm [50] was used with a Poisson model describing the relationship between the nodes. The number of blocks was chosen using a penalized likelihood criterion (ICL), and the likelihood maximization was obtained via a variational version of the Expectation-Maximization algorithm.

### Statistical analysis and figure construction

To compare WMA or the prevalence among studies of ASVs belonging to different groups (taxonomic rank or SBM cluster), the non-parametric Wilcoxon-Mann-Whitney test was performed. To create figures, the R packages ggplot2, viridis, ggpubr, and ComplexHeatmap were used [51, 52].

## ACKNOWLEDGMENTS

We are grateful to INRAE MICA Division and the ENS Paris-Saclay for the funding of this PhD and to INRAE MIGALE bioinformatics facility (MIGALE, INRAE, 2020. Migale bioinformatics Facility, doi:10.15454/1.5572390655343293E12) for providing computing and storage resources.

## DATA AVAILABILITY STATEMENT

No new research data was generated in the preparation of this article.

